# Long-term sterile immunity induced by an adjuvant-containing live-attenuated AIDS virus

**DOI:** 10.1101/2020.05.22.111781

**Authors:** Tomotaka Okamura, Yuya Shimizu, Tomohiro Kanuma, Yusuke Tsujimura, Masamitsu N Asaka, Kazuhiro Matsuo, Takuya Yamamoto, Yasuhiro Yasutomi

## Abstract

Antigen 85B (Ag85B) is one of the most dominant proteins secreted from most mycobacterial species, and it induces Th1-type immune responses as an adjuvant. We genetically constructed a live attenuated simian human immunodeficiency virus to express the adjuvant molecule Ag85B (SHIV-Ag85B). SHIV-Ag85B could not be detected 4 weeks after injection in cynomolgus macaques, and strong SHIV-specific T cell responses were induced in these macaques. When these macaques in which SHIV-Ag85B had become undetectable were challenged with pathogenic SHIV89.6P at 37 weeks after SHIV-Ag85B became undetectable, SHIV89.6P could not be detected after the challenge. Eradication of SHIV89.6P was confirmed by adoptive transfer experiments and CD8-depletion studies. The SHIV-Ag85B-inoculated macaques showed enhancement of Gag-specific monofunctional and polyfunctional CD8^+^ T cells in the acute phase of pathogenic SHIV challenge. The results suggest that SHIV-Ag85B elicited strong sterile immune responses against pathogenic SHIV and that it may lead to the development of a vaccine for AIDS virus infection.

**Importance:** Development of an effective HIV vaccine has been a major priority to control the worldwide AIDS epidemic. The moderately attenuated prototypic vaccine strain SIVmac239Δnef has been used in various studies; however, it does not provide sufficient effects to prevent infection. The use of adjuvant in vaccination is thought to be useful for enhancing the immune responses to various pathogens. In the present study, we constructed a live attenuated SHIV virus expressing adjuvant molecule Ag85B and assessed vaccine effects in cynomolgus macaques. The present study shows that live-attenuated SHIV expressing Ag85B elicits viral antigen-specific polyfunctional CD8^+^ T cell responses against pathogenic SHIV and provide the possibility of eradicating a pathogenic lentivirus from infected animals.

## Introduction

Despite the considerable resources that have been committed to developing an effective HIV vaccine over the past three decades, this objective remains elusive. Although anti-retroviral therapy has led to a dramatic reduction in HIV-related morbidity and mortality, it is not a cure (1). Recently, there have been two cases in which HIV remission was achieved by cell transplantation (2, 3). The development of an effective vaccine could prevent the spread of HIV infection, but vaccination efforts have been relatively unsuccessful over the past three decades (4, 5). The only successful HIV vaccine to date, evaluated in the RV144 clinical trial, showed an overall efficacy of only 31% (6).

Various vaccine viruses attenuated by genetic disruption of key regulatory genes including *nef, vpx, vpr*, and *vif*, have been used in previous studies, though the moderately attenuated prototypic vaccine strain SIVmac239Δnef has been used in most studies (7-11). The live attenuated immunodeficiency viruses *nef*-deleted SIV and SHIV have proven to be highly effective for vaccines in non-human primate models, but they are not sufficiently safe to use as a template for HIV vaccines in humans (12-14). One promising strategy for improving the immunodeficiency of a live attenuated virus is to genetically engineer the virus to co-express an immunostimulatory agent such as a cytokine or a chemokine. Several studies have demonstrated that insertion of the cytokine or chemokine gene in live attenuated and genetically defective SIV or SHIV can improve the immunogenicity and enhance the protective capacity of the virus (15-17) compared to those of safer, less virulent strains.

Antigen 85B (Ag85B) is considered to be an immunogenic protein that can induce a strong T helper type 1 (Th1) immune response in hosts sensitized by BCG. Ag85B, which belongs to the Ag85 family, is one of the most dominant proteins secreted from most mycobacterial species. It has been shown to induce substantial Th1 cell proliferation and vigorous Th1 cytokine production in mice (18). Ag85B has been reported to induce Th1 cells in immunotherapy for atopic dermatitis and allergic asthma (19-21) and in tuberculosis vaccination (22). These results make Ag85B an attractive candidate for an immune adjuvant.

To study the adjuvant effect of Ag85B in SHIV macaque models, cDNA of *Ag85B* was inserted into the *nef* gene-eliminated site of SHIV, generating SHIV-Ag85B. We then analyzed immune responses in cynomolgus macaques inoculated with SHIV-Ag85B and with parental SHIV-NI and also examined the long-term protective efficacy in those macaques after challenge with pathogenic SHIV89.6P.

## Results

### Construction of SHIV-Ag85B

A recombinant SHIV was engineered to express Ag85B in place of *nef* in SHIV-NI (Fig. S1A). Expression of Ag85B was detected by Western blot analysis using the cell lysate from a human lymphoid cell line (M8166) infected with SHIV-Ag85B (Fig. S1B). SHIV-ag85B replicated well not only in a human lymphoid cell line (CEM×174) but also in a macaque lymphoid cell line (HSC-F) (Fig. S1C). The replication profile of SHIV-Ag85B in lymphoid cell lines was similar to that of parental SHIV-NI.

### Direct effects of SHIV-Ag85B infection in cells *in vitro*

We next examined direct effects of SHIV-Ag85B infection such as innate immune responses. Immune responses were assessed at 48 h after infection with SHIV-Ag85B in CEM×174 cells. Expression levels of TNF-α after infection with SHIV-Ag85B were significantly higher than those after infection with SHIV-NI. In contrast, mRNA levels of IFN-γ, IL-6, and type I IFNs were almost the same after infection with SHIV-Ag85B and after infection with SHIV-NI (Fig. S1D). Double-strande RNA (dsRNA) is a dominant activator of innate immunity because viral dsRNA is recognized by RIG-I and MDA5 (23, 24). The mRNA expression of the intracellular receptors RIG-I and MDA5 was also enhanced by infection with SHIV-Ag85B (Fig. S1E). LGP2 mRNA levels induced by SHIV-Ag85B were similar to those induced by SHIV-NI.

### Viral loads in cynomolgus macaques inoculated with SHIV-Ag85B

To study the adjuvant effect of Ag85B in SHIV macaque models, two experiments were performed at different times. The design of the macaque study is outlined in Fig. 1A. In the first experiment (Exp 1), all macaques exhibited peak viremia about 2 to 3 weeks after inoculation (Fig. 1B). The plasma viral loads in macaques inoculated with SHIV-Ag85B declined earlier than those in macaques inoculated with SHIV-NI. The plasma viral load in macaques inoculated with SHIV-Ag85B fell below the measurable limit at 2 to 4 weeks after inoculation, whereas that in macaques inoculated with SHIV-NI reached an undetectable level at about 8 to 12 weeks after inoculation (Fig. 1B). In contrast, the plasma viral loads in macaques inoculated with SHIV89.6P were maintained at high levels after the acute phase of inoculation (Fig. 1B). Due to limited viral replication, proviral DNAs in SHIV-Ag85B-inoculated macaques were detected up to 4 weeks after inoculation and became undetectable in SHIV-Ag85B-inoculated macaques at 8 weeks after inoculation (Fig. 1D). The stability of the inserted *Ag85B* gene in SHIV-Ag85B was analyzed by PCR. The full length of the inserted *Ag85B* gene in PBMCs from macaques inoculated with SHIV-Ag85B was detected 2 weeks after inoculation (Fig. S2). SHIV-NI-inoculated and SHIV89.6P-inoculated macaques showed proviral DNAs in PBMCs examined at all stages of infection (Fig. 1D). After the inoculation, peripheral blood CD4^+^ T cells remained within the normal range of levels in both SHIV-Ag85B- and SHIV-NI-inoculated macaques, and these macaques remained healthy clinically (Fig. 1C). In contrast, SHIV89.6P-inoculated macaques showed very low CD4^+^ T cell counts (< 150 cells/μl) during the observation period (Fig. 1C). In the second experiment (Exp 2), all of the macaques inoculated with SHIV-Ag85B showed viremia within 2 weeks after inoculation (Fig. 1B). In these macaques, plasma viral RNAs and proviral DNAs were reduced to almost undetectable levels within 4 to 8 weeks after inoculation and the numbers of peripheral CD4^+^ T cells were maintained at normal levels (Fig. 1B to D). SHIV-Ag85B infection was confirmed in all experimental animals for at least up to 4 weeks after inoculation; however, SHIV-Ag85B might have been eradicated from macaques after 8 weeks of inoculation.

**FIG 1.**
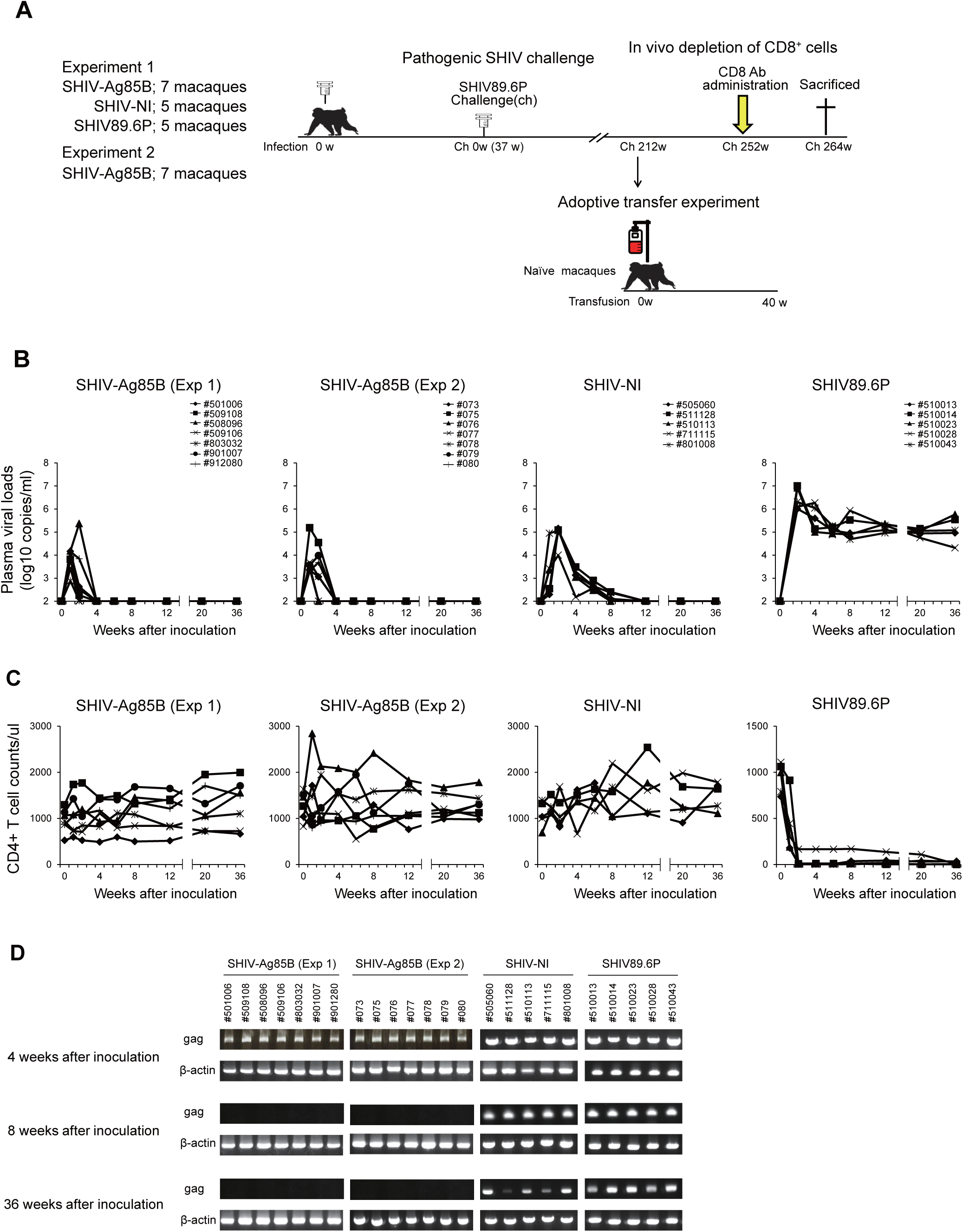
Kinetics of viral loads and detection of proviral DNA in macaques inoculated with SHIVs. (A) Experimental design, SHIV infection, adoptive transfer experiment, CD8^+^ cell depletion study and necropsy time points. (B) Plasma viral loads in macaques inoculated with SHIV-Ag85B (Exp 1), SHIV-Ag85B (Exp 2), SHIV-NI and SHIV89.6P. Plasma viral loads were measured by quantitative RT-PCR. The detection limit of plasma viral load was 100 copies/ml. (C) Proviral DNAs in PBMCs of the macaques after SHIV-Ag85B (Exp 1), SHIV-Ag85B (Exp 2), SHIV-NI and SHIV89.6P inoculation were detected by nested PCR for the SIV *gag* specific region. β-actin was used as control. (D) Changes in the percentages of CD4^+^ T cells after SHIV-Ag85B (Exp 1), SHIV-Ag85B (Exp 2), SHIV-NI and SHIV89.6P infection in cynomolgus macaques. Whole blood was stained by CD3, CD4 and CD8 Abs, and CD4^+^ T cell counts were determined by flow cytometric analysis.

### SHIV-Ag85B-induced antigen-specific CD8^+^ immune response

SHIV antigen (Gag/pol)-specific T cell responses were measured by an IFN-γ ELISPOT assay of PBMCs. The IFN-γ ELISPOT responses to Gag/pol in SHIV-Ag85B-inoculated macaques at 2 weeks after inoculation were stronger than those in other groups of animals, and then the responses were decreased at 8 weeks after inoculation (Fig. 2A). We also analyzed Ag85B-specific IFN-γ ELISPOT responses of PBMCs from SHIV-Ag85B-inoculated macaques at 2 weeks after inoculation. These macaques showed weak immune responses to Ag85B in ELISPOT analysis (Fig. S3). IFN-γ ELISPOT responses to Gag/pol were detected in SHIV-NI-inoculated macaques at 2 weeks and were slightly increased at 8 weeks after inoculation. The macaques inoculated with SHIV89.6P showed weak immune responses at all time points in ELISPOT analysis (Fig. 2A). To determine whether polyfunctional Gag/pol-specific CD8^+^ T cell immune responses were induced in SHIV-Ag85B-inoculated macaques, we analyzed intracellular cytokine staining for IFN-γ, TNF-α, and IL2 by flow cytometry. SHIV-Ag85B-inoculated macaques showed preferential accumulation of Gag/pol-specific IFN-γ or TNF-α-single-positive and IFN-γ and TNF-α-double-positive CD8^+^ T cells in PBMCs at 2 weeks after inoculation, and these responses were decreased at 8 weeks after inoculation (Fig. 2B). SHIV-Ag85B also elicited antigen-specific CD4^+^ T cells producing a single cytokine (Fig. S4). In the SHIV-NI-inoculated macaques, the percentages of antigen-specific CD8^+^ T cells and CD4^+^ T cells were lower than those in SHIV-Ag85B-inoculated macaques at 2 weeks after inoculation; however, the percentages of those cells in SHIV-NI-inoculated macaques were higher than those in SHIV-Ag85B-inoculated macaques at 8 weeks after inoculation. In the SHIV89.6P-inoculated macaques, the percentages of those cells were lower at all time points (Fig. 2B and Fig. S4).

**FIG 2.**
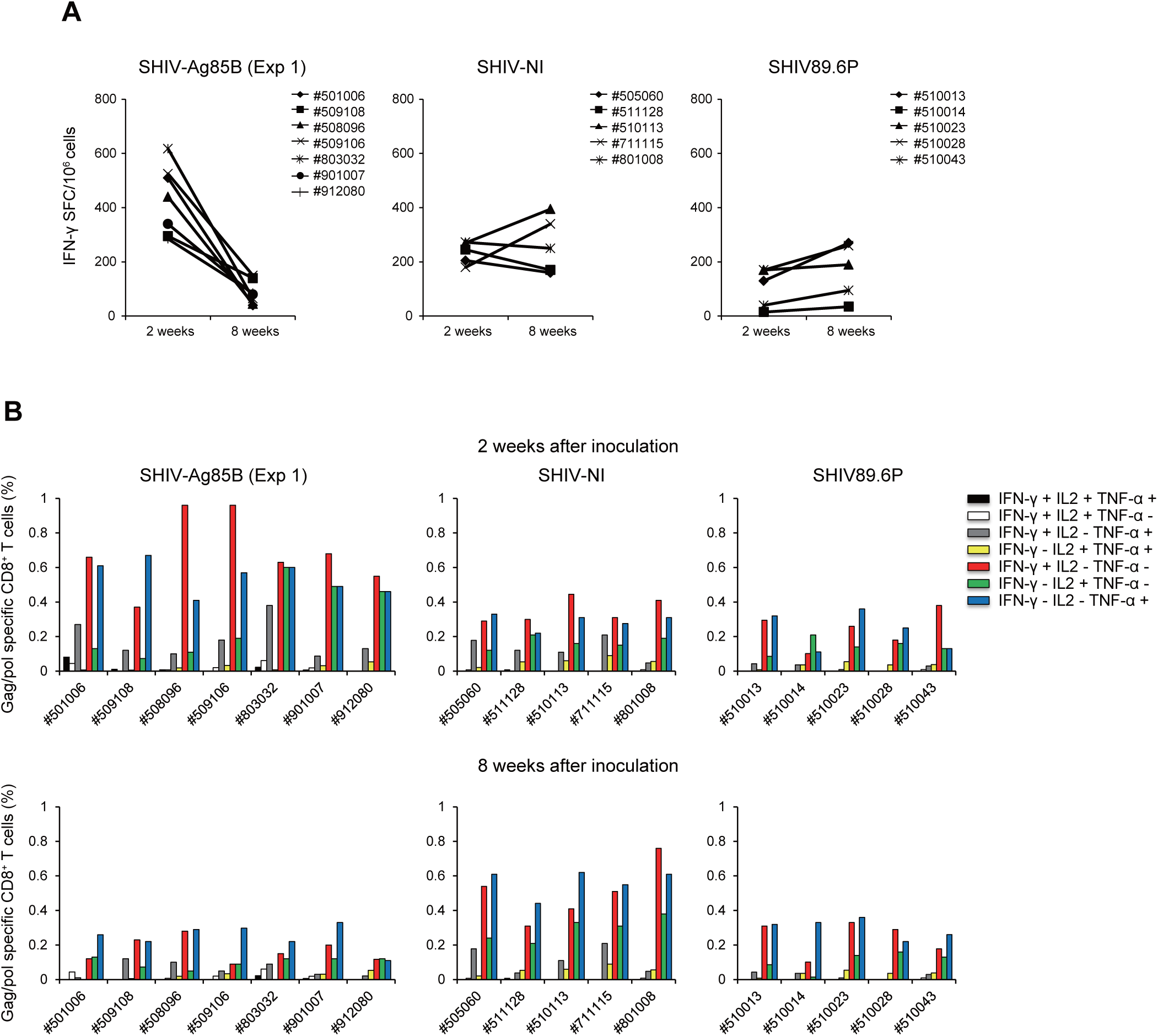
SHIV antigen-specific T cell responses in macaques inoculated with SHIVs. (A) Numbers of Gag/pol-specific IFN-γ-producing cells in macaques inoculated with SHIV-Ag85B (Exp 1), SHIV-NI and SHIV89.6P were determined by ELISPOT assays. PBMCs obtained at 2 weeks and 8 weeks after SHIV inoculation were co-cultured for 6 h with autologous B-LCL cells that had been infected with a recombinant vaccinia virus expressing SIV Gag/pol. Antigen-specific IFN-γ ELISPOT results are depicted as spots per 10^6^ PBMCs. (B) Percentage of Gag/pol-specific CD8^+^ T cells producing IFN-γ, TNF-α, or IL2 in macaques inoculated with SHIV-Ag85B (Exp 1), SHIV-NI and SHIV89.6P. The cytokine profile in cells was determined by flow cytometry by gating for lymphocytes and CD8^+^ T cells. PBMCs obtained at 2 weeks and 8 weeks after SHIV inoculation were co-cultured for 6 h with autologous B-LCL cells that had been infected with a recombinant vaccinia virus expressing SIV Gag/pol.

SHIV-specific antibody titers in plasma of SHIV-Ag85B, SHIV-NI and SHIV89.6P-inoculated macaques were examined by using a passive agglutination method. All of the macaques showed antibody responses against SHIV at 4 to 8 weeks after inoculation. The antibody responses increased in SHIV-NI-inoculated macaques at later time points, whereas the responses were marginal in SHIV-Ag85B-inoculated macaques (Fig. S5).

### Challenge with a pathogenic SHIV89.6P

To know the effect of pathogenic virus infection in SHIV-Ag85B-undetected macaques or SHIV-NI-infected macaques, all of the macaques were challenged intravenously with 10^5^ TCID_50_ of heterologous pathogenic SHIV89.6P at 37 weeks after inoculation. The peak plasma viral loads in SHIV-Ag85B-undetected macaques were similar to those in SHIV-NI-infected macaques (Fig. 3A). The plasma viral loads in six of the seven SHIV-Ag85B-undetected macaques declined to almost below the detection level from 6 to 12 weeks after the challenge; however, viral RNAs remained negative for more than 250 weeks (in the chronic phase) in six macaques. Moreover, CD4^+^ T cells in those macaques were maintained at the normal level (Fig. 3B). In addition, six of the SHIV-Ag85B-undetected macaques did not show progression to obvious AIDS-like disease. In the second experiment, six of the seven SHIV-Ag85B-undetected macaques did not show any detectable plasma viral load from 6 to 16 weeks after the challenge (Fig. 3A). In the chronic phase, viral RNAs remained negative for more than 20 weeks in six macaques. The number of CD4^+^ T cells was maintained at the normal level in all of these macaques (Fig. 3B). In contrast, the plasma viral loads of two SHIV-Ag85B-undetected macaques (#509108 and #078) were similar to those of SHIV-NI-infected macaques. CD4^+^ T cells in one macaque (#509108) gradually decreased during the observation period. That macaque (#509108) showed AIDS symptoms at 989 days after the challenge. In the SHIV-NI-infected macaques, the set-point plasma viral loads were less than 10^4^ copies/ml after the acute phase (Fig. 3A). CD4^+^ T cells in these macaques gradually decreased during the observation period (Fig. 3B). Thereafter, two of the SHIV-NI-infected macaques (#510113 and #801008) showed AIDS symptoms at 718 days and 1450 days after the challenge, respectively. In the control macaques, viremia was maintained at high levels during the observation period (Fig. 3A). These macaques showed very low CD4^+^ T cell counts (< 100 cells/μl) during the observation period (Fig. 3B) and showed AIDS symptoms at 402 to 1225 days after the challenge.

**FIG 3.**
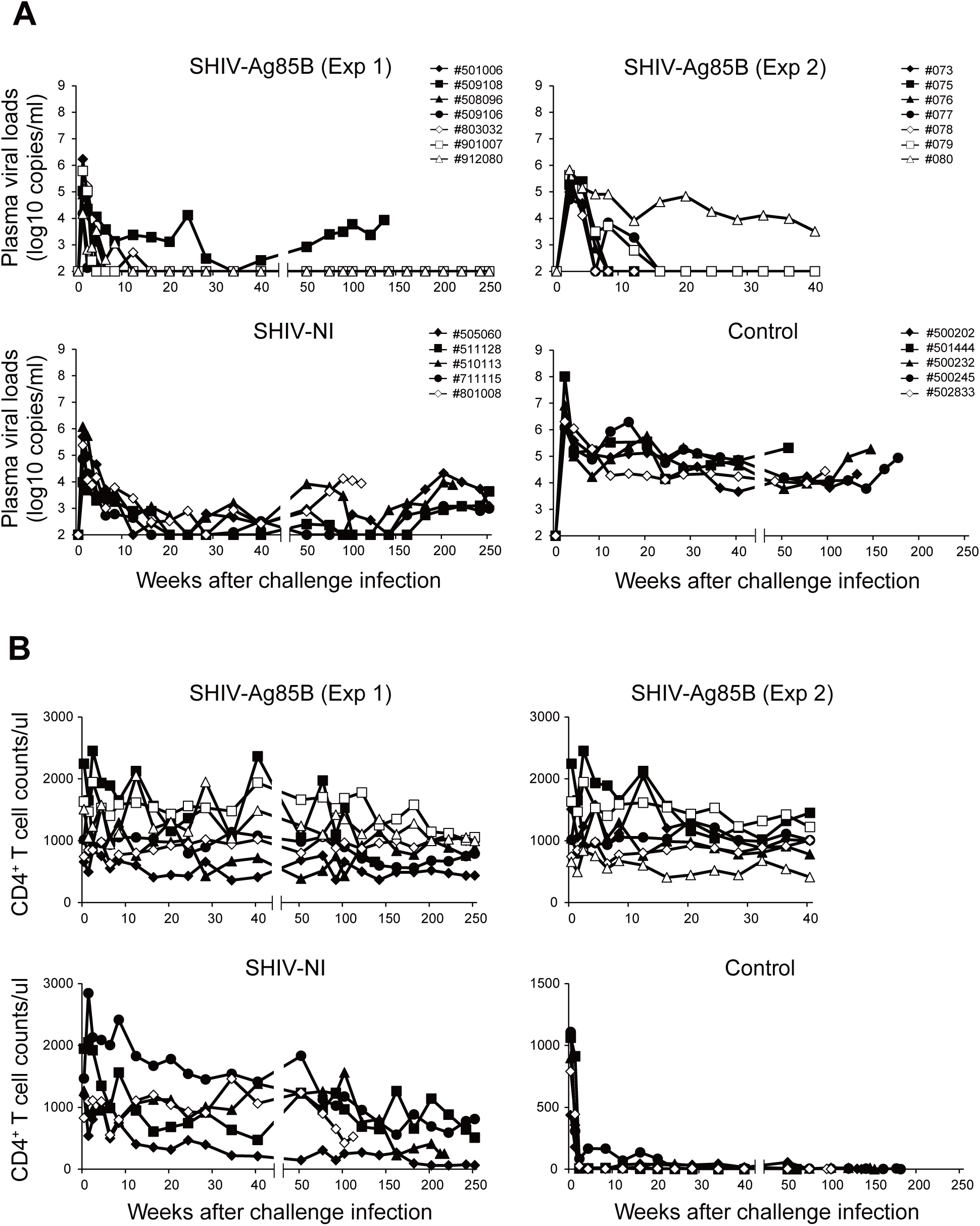
Kinetics of viral loads, CD4^+^ T cells and proviral DNA in macaques inoculated with SHIV-Ag85B or SHIV-NI after pathogenic SHIV89.6P challenge. (A) Plasma viral loads in macaques inoculated with SHIV-Ag85B (Exp 1), SHIV-Ag85B (Exp 2), and SHIV-NI after SHIV89.6P challenge. As controls, 5 naïve macaques were challenged intravenously with SHIV89.6P. Plasma viral loads were measured by quantitative RT-PCR. The detection limit of plasma viral load was 100 copies/ml. (B) Count of CD4^+^ T cells was analyzed by flow cytometry and whole blood cell counts. Whole blood was stained by CD3, CD4 and CD8 Abs, and CD4^+^ T cell counts were determined by flow cytometric analysis.

### Quantification of proviral DNA after pathogenic SHIV89.6P challenge

To better understand the viral status in SHIV-Ag85B-undetected macaques (#501006, #509106, #803032, #901007, #508096, #912080 and #509108), proviral DNAs in PBMCs and lymphoid tissues after SHIV89.6P challenge were measured by ultrasensitive digital PCR. All SHIV-Ag85B-undetected macaques showed proviral DNA loads in PBMCs at 12 or 24 weeks after the challenge (Fig. 4A). Strikingly, in a period of 36 weeks after the challenge, four of the SHIV-Ag85B-undetected macaques (#501006, #509106, #803032 and #901007) did not exhibit any proviral DNA of SHIV in PBMCs, and proviral DNA in PBMCs remained negative for more than 50 weeks in four macaques (Fig. 4A). Moreover, proviral DNA was not detected in lymphoid tissues of those macaques (Fig. 4B). Also, SIV Nef-positive cells were not detected by immunohistochemistry in lymphoid tissues from four macaques (Fig. 4C). Two SHIV-Ag85B-undetected macaques (#508096 and #912080) without a plasma viral load showed proviral DNAs in PBMCs and lymphoid tissues during the observation period (Fig. 4A and B). Control macaques and the SHIV-NI-infected macaques showed high levels of proviral DNAs in PBMCs and lymphoid tissues after SHIV-89.6P challenge (Fig. 4A and B).

**FIG 4.**
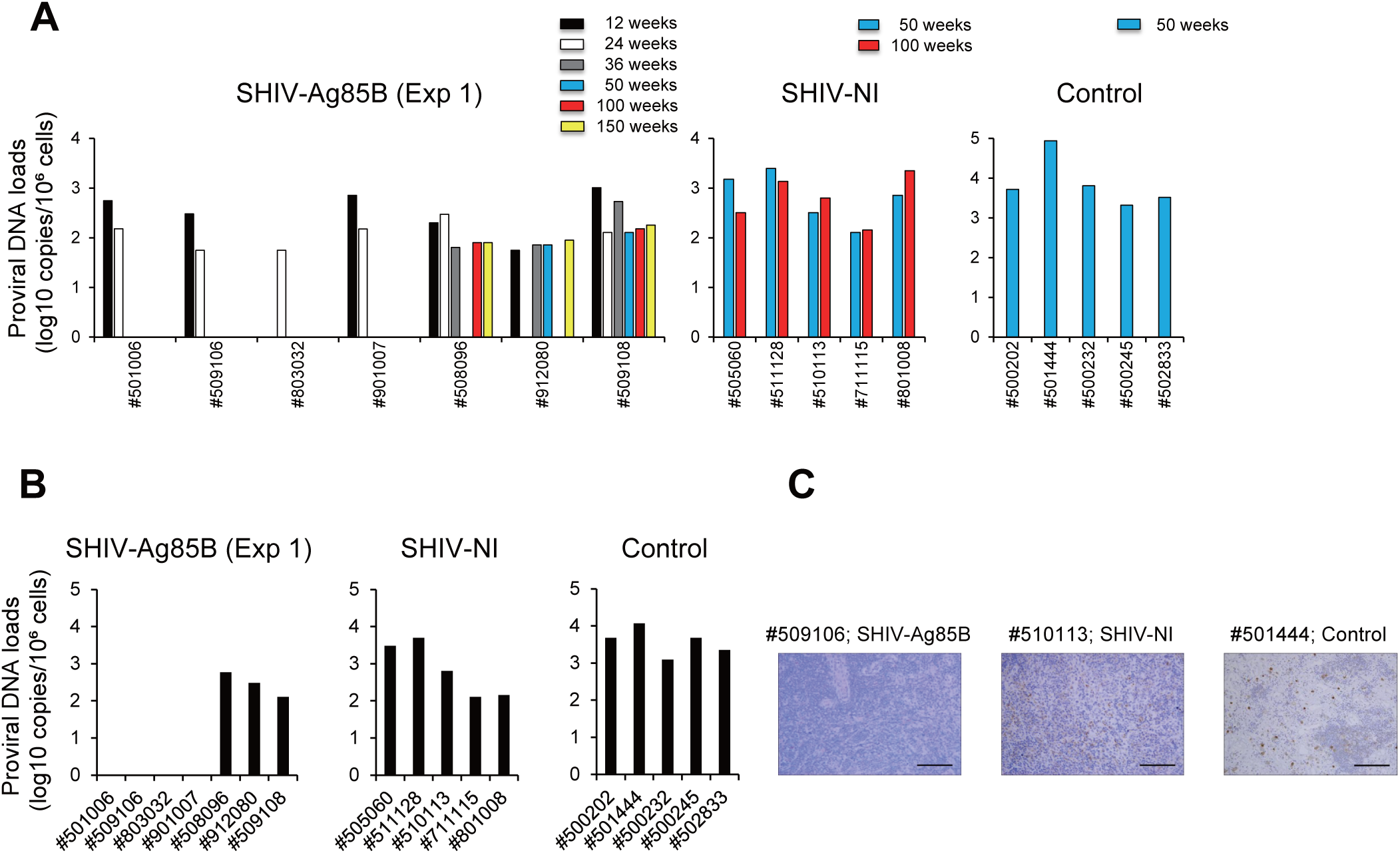
Kinetics of proviral DNA in macaques inoculated with SHIV-Ag85B or SHIV-NI after pathogenic SHIV89.6P challenge. (A) Proviral DNA loads in PBMCs of SHIV-Ag85B-undetected macaques (Exp 1) or SHIV-NI-infected macaques after SHIV89.6P challenge as measured by ultrasensitive digital PCR. The detection limit of proviral DNA load was 5 copies per 10^6^ cells. (B) Proviral DNA loads in lymphoid tissues of SHIV-Ag85B-undetected macaques (Exp 1) or SHIV-NI-infected macaques at 34 weeks after the challenge as measured by ultrasensitive digital PCR. (C) Immunohistochemical detection of SHIV Nef antigen in inguinal lymph nodes at 34 weeks after the challenge. Brown staining indicates SHIV Nef-positive cells. Magnification×200. Black scale bars, 100 μm.

### Protective antigen-specific immune responses after pathogenic SHIV89.6P challenge

Four of the seven SHIV-Ag85B-inoculated macaques (#501006, #509106, #803032 and #901007) showed no increase in overall plasma viral RNA and proviral DNA, and small amounts of proviral DNA were detected in two macaques (#508096 and #912080) in the SHIV-Ag85B-inoculated macaques. One SHIV-Ag85B-inoculated macaque (#509108) showed a low level of plasma viral RNA that was similar to the levels in five SHIV-NI-inoculated macaques (#505060, #511128, #510113, #711115 and #801008). Those 6 macaques (one SHIV-Ag85B-inoculated macaque and five SHIV-NI-inoculated macaques) showed low levels of plasma viral RNA compared to the levels in control macaques, and five SHIV89.6P-injected naïve macaques (#500202, #501444, #500232, #500245 and #502833) showed stable viremia above 10^4^ to 10^7^ copies/ml of viral RNA (Fig. 3A and 4A). We next analyzed SHIV antigen-specific production of IFN-γ by ELISPOT assays in these experimental macaques after the challenge. In six SHIV-Ag85B-undetected macaques, ELISPOT responses were increased dramatically at 2 weeks after the challenge (Fig. 5A). The responses were reduced in most of those macaques at 12 weeks after the challenge. In only one of the SHIV-Ag85B-undetected macaques, the IFN-γ ELISPOT response was moderately increased at 2 weeks after the challenge and was maintained at 12 weeks after the challenge. The response was similar to ELISPOT responses of SHIV-NI. The control macaques showed weak IFN-γ ELISPOT responses at all time points (Fig. 5A). The polyfunctionality of Gag/pol-specific T cells was also examined by flow cytometry. On the day of the challenge, we did not detect polyfunctional Gag/pol-specific CD4^+^ and CD8^+^ T cells in any of the macaques (Fig. 5B and Fig. S6). In six SHIV-Ag85B-undetected macaques, there was preferential accumulation of Gag/pol-specific, triple-positive (IFN-γ, TNF-α and IL2), double-positive (IFN-γ and TNF-α), and IFN-γ or TNF-α-single positive CD8^+^ T cells at 2 weeks after the challenge (Fig. 5B). In addition, the percentages of these polyfunctional Gag/pol-specific CD8^+^ T cells in six SHIV-Ag85B-undetected macaques were higher than those in SHIV-NI-infected macaques and control macaques (Fig. 5B). We found polyfunctional Gag/pol-specific CD4^+^ T cells in six SHIV-Ag85B-undetected macaques, but the percentages were similar to those in SHIV-NI-infected macaques at 2 weeks after the challenge (Fig. S6). We also analyzed the memory phenotype of SHIV antigen-specific CD8^+^ T cells in those three groups. The percentages of Gag/pol-specific IFN-γ-producing CD8^+^ T cells in six SHIV-Ag85B-undetected macaques were higher than those in SHIV-NI-infected macaques and control macaques (Fig. S7). In those macaques, Gag/pol-specific responses were a mixed effector (EM) and central memory (CM) phenotype for CD8^+^ T cells (Fig. S7).

**FIG 5.**
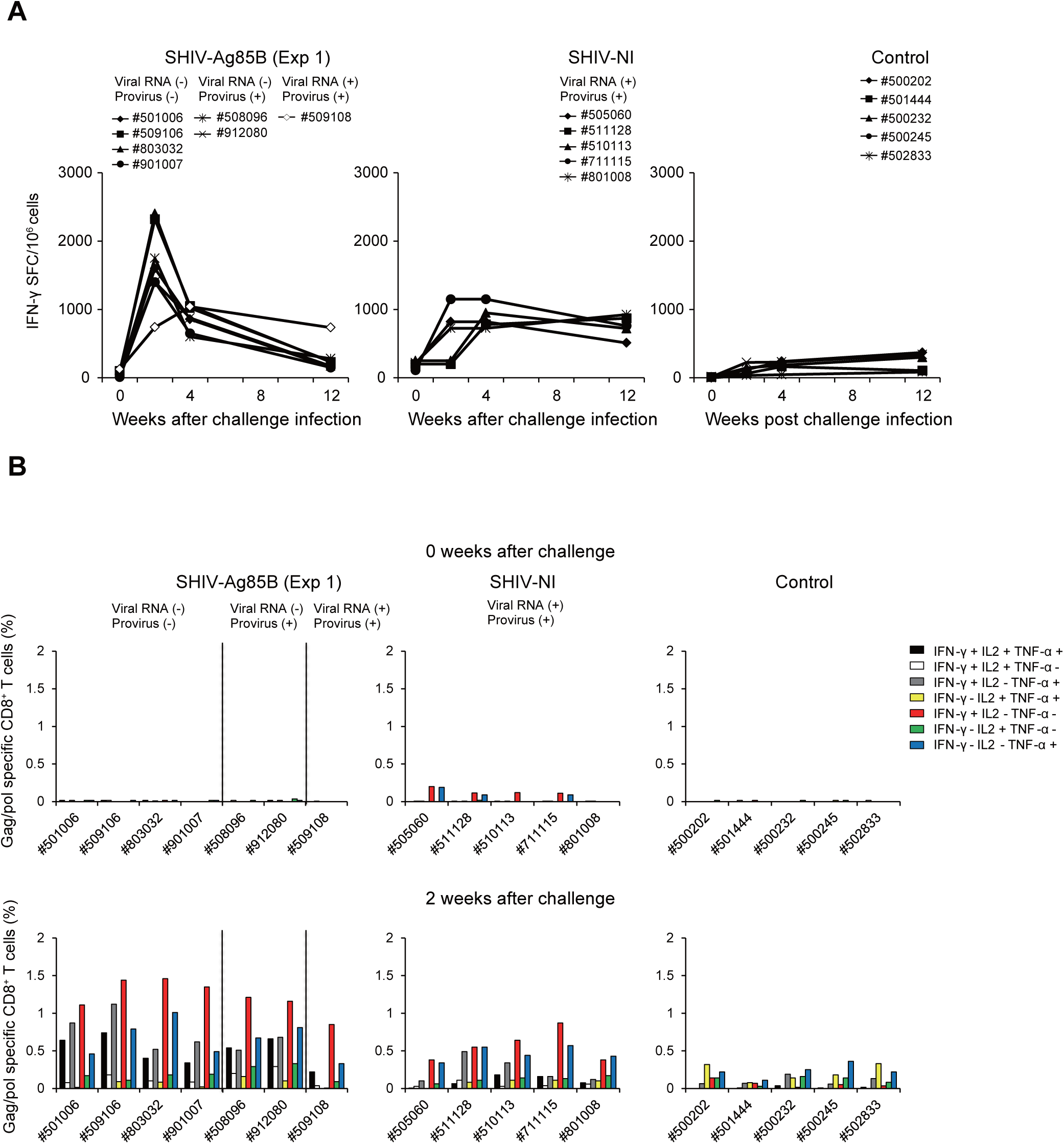
Protective antigen-specific T cell responses after pathogenic SHIV89.6P challenge. (A) Kinetics of Gag/pol-specific IFN-γ-producing cells in SHIV-Ag85B (Exp 1)-undetected macaques, SHIV-NI-infected macaques and control macaques after SHIV89.6P challenge were measured by ELISPOT assays. PBMCs obtained at 0 weeks, 2 weeks, 4 weeks, and 12 weeks after the challenge were co-cultured for 9 h with autologous B-LCL cells that had been infected with a recombinant vaccinia virus expressing SIV Gag/pol. Antigen-specific IFN-γ ELISPOT results are depicted as spots per 10^6^ PBMCs. (B) Percentages of Gag/pol-specific CD8^+^ T cells producing IFN-γ, TNF-α, or IL2 in SHIV-Ag85B (Exp 1)-undetected macaques, SHIV-NI-infected macaques and control macaques after SHIV89.6P challenge. The cytokine profile in cells was determined by flow cytometry by gating for lymphocytes and CD8^+^ T cells. PBMCs obtained at 0 weeks and 2 weeks after the challenge were co-cultured for 6 h with autologous B-LCL cells that had been infected with a recombinant vaccinia virus expressing SIV Gag/pol.

After the pathogenic SHIV89.6P challenge, we measured SHIV-specific antibody in plasma of SHIV-Ag85B-undetected macaques, SHIV-NI-infected macaques and control macaques. Antibody titers in SHIV-Ag85B-undetected macaques and SHIV-NI-infected macaques rapidly increased and reached peak levels within 4 weeks after the challenge (Fig. S8). In the six SHIV-Ag85B-undeteced macaques, anti-SHIV antibody titers then gradually decreased during the observation period. In contrast, one SHIV-Ag85B-undetected macaque (#509108) and five SHIV-NI-inoculated macaques maintained strong antibody responses (Fig. S8).

### Eradication of pathogenic SHIV89.6P in SHIV-Ag85B-undetected macaques

To examine the eradication of virus in the animals, we performed adoptive transfer experiments and infused peripheral blood and lymph node mononuclear cells by the intravenous route from six SHIV-Ag85B-undetected macaques (#501006, #508096, #509106, #803032, #901007, and #912080) and three SHIV-NI-infected macaques (#505060, #511128 and #711115) into naïve cynomolgus macaques (Fig. 6A). Adoptive transfer of cells from all SHIV-NI-infected macaques at 212 weeks after the challenge readily transferred typical SHIV infection to naïve macaques. In addition, adoptive transfer of cells from two SHIV-Ag85B-undetected macaques (#508096 and #912080) at 212 weeks after the challenge readily transferred infection to naïve macaques and the macaques showed plasma viral RNA and proviral DNA, indicating the presence of replicating virus in those macaques (Fig. 6B and C). In contrast, adoptive transfer of cells from four SHIV-Ag85B-undetected macaques (#501006, #509106, #803032, and #901007) at 212 weeks after the challenge did not transfer SHIV infection to naïve macaques (Fig. 6B and C), indicating that cells containing pathogenic virus were not present in these cell preparations from SHIV-Ag85B and SHIV-89.6P-undetected macaques.

**FIG 6.**
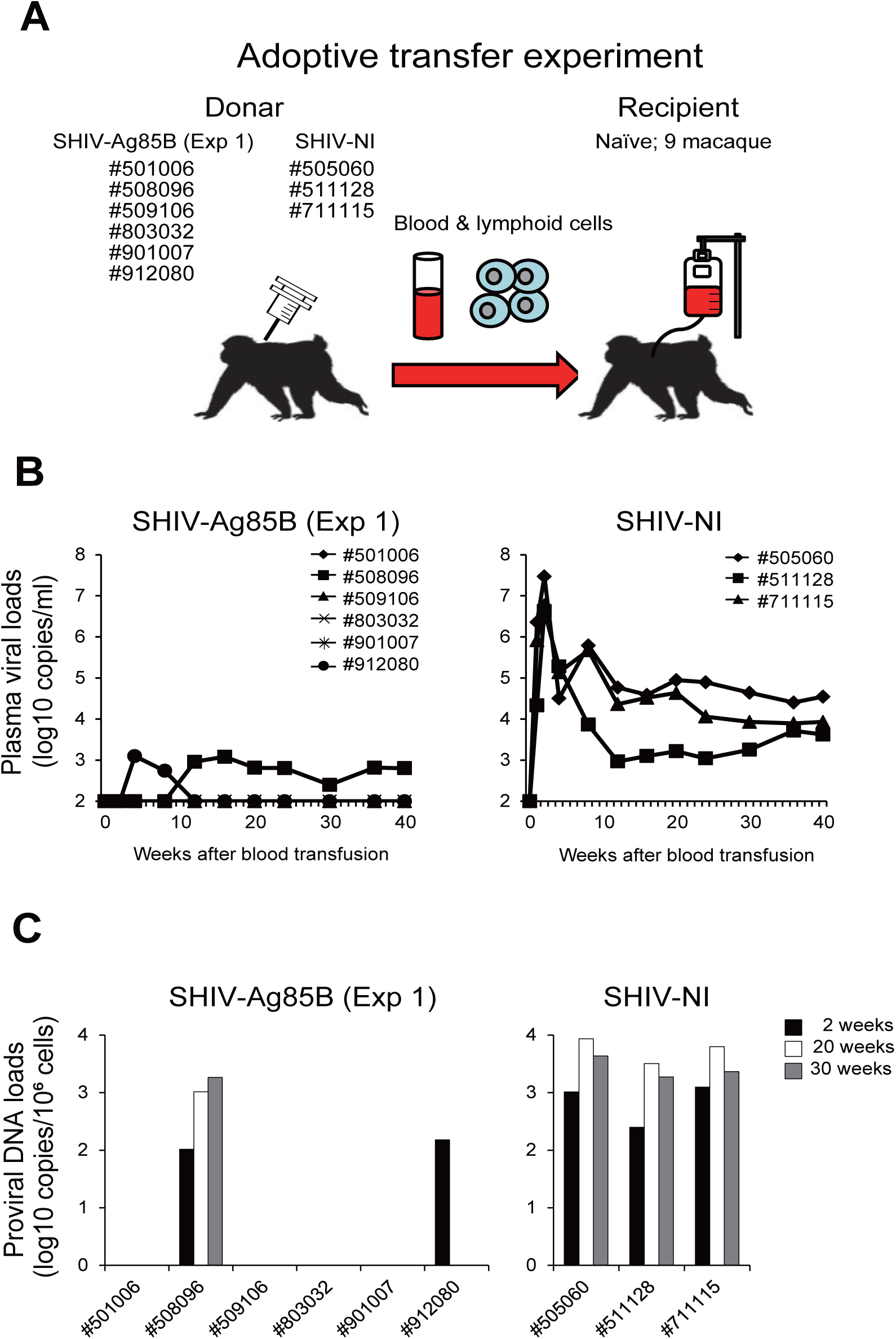
Kinetics of viral loads and proviral DNA after the adoptive transfer experiment. (A) Schematic of the adoptive transfer experiment. (B) Plasma viral loads following adoptive transfer are shown. Plasma viral loads were measured by quantitative RT-PCR. The detection limit of plasma viral load was 100 copies/ml. (C) Proviral DNA loads in PBMCs following adoptive transfer are shown. Proviral DNA loads were measured by ultrasensitive digital PCR. The detection limit of proviral DNA load was 5 copies per 10^6^ cells.

We next depleted CD8^+^ cells in six SHIV-Ag85B-undetected macaques (#509108, #505060, #511128, #510113, #711115 and #801008) and three SHIV-NI-infected macaques (#505060, #511128 and #711115) at 252 weeks after the SHIV89.6P challenge. After infusion, all of the macaques exhibited a depletion of CD8^+^ cells in peripheral blood (Fig. 7A). In the absence of CD8^+^ cells, two SHIV-Ag85B-undetected macaques (#508096 and #912080) showed a rapid spike in plasma viral RNA, which subsequently declined to an undetectable level (Fig. 7B). On the other hand, plasma viral loads in all of the SHIV-NI-infected macaques were maintained at 10^4^ copies/ml of viral RNA after administration. One macaque (#511128) showed symptoms of AIDS at 2 weeks after administration. In contrast, in four SHIV-Ag85B-undetected macaques (#501006, #509106, #803032 and #901007), plasma viral loads were not detected at any time point after CD8^+^ cell depletion (Fig. 7B).

**FIG 7.**
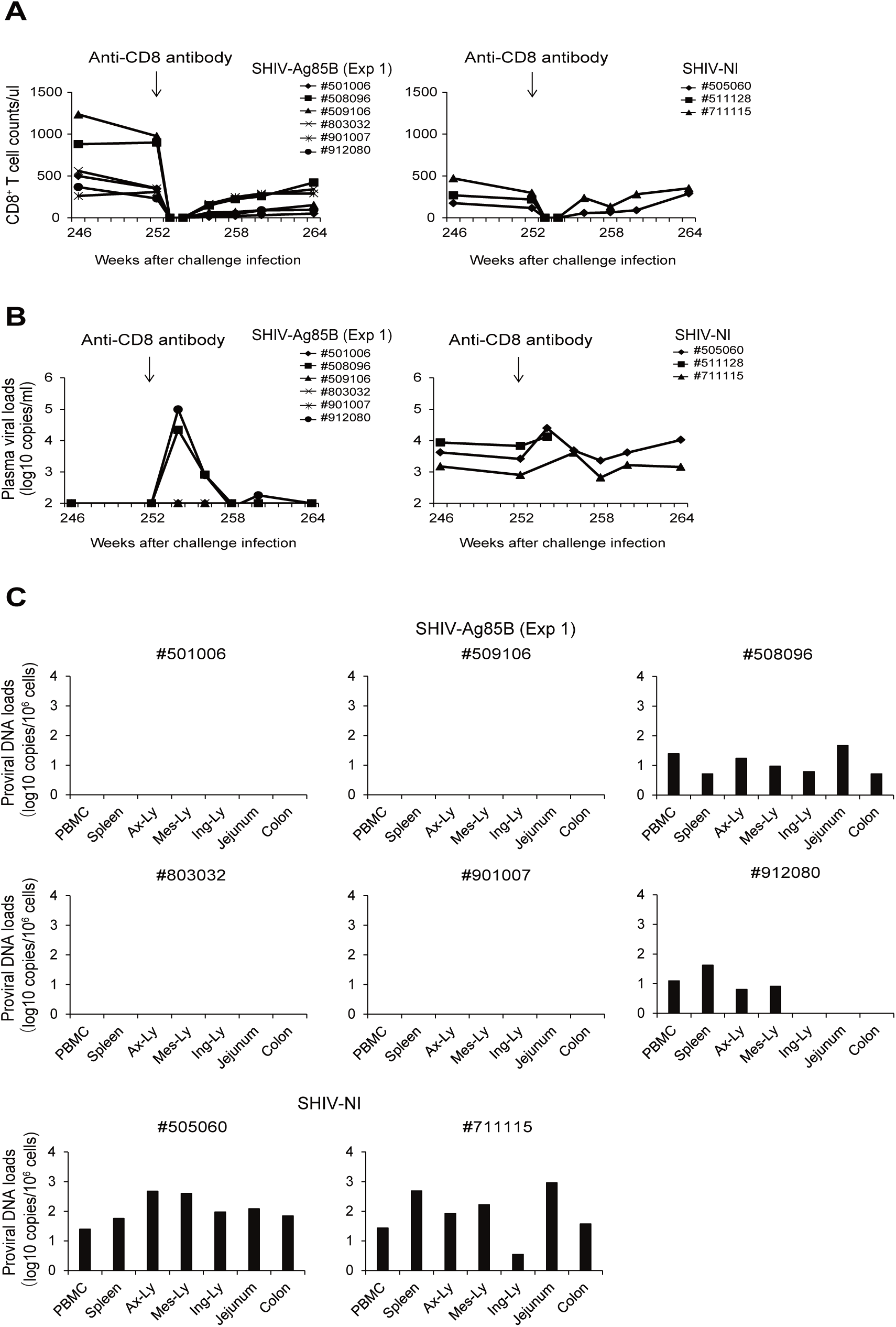
Virology analysis in SHIV-Ag85B-mediated protection following CD8^+^ cell depletion. (A) Changes in peripheral CD8^+^ T cell counts after anti-CD8 antibody administration. SHIV-Ag85B (Exp 1)-undetected macaques and SHIV-NI-infected macaques were administered an anti-CD8^+^ antibody at a dose of 15 mg per kg body weight at 252 weeks after SHIV89.6P challenge. (B) Kinetics of plasma viral RNA loads after administration of the anti-CD8 antibody. Plasma viral loads were measured by quantitative RT-PCR. The detection limit of plasma viral load was 100 copies/ml. (C) At necropsy, SHIV proviral DNA loads in PBMCs, the spleen, axillary lymph node (Ax-Ly), mesenteric lymph node (Mes-Ly), inguinal lymph node (Ing-Ly), jejunum and colon of SHIV-Ag85B (Exp 1)-undetected macaques and SHIV-NI-infected macaques were measured by ultrasensitive digital PCR. The detection limit of proviral DNA load was 5 copies per 10^6^ cells.

To examine virus clearance in the animals, proviral DNAs in various lymphoid tissues of SHIV-Ag85B-undetected macaques were measured by ultrasensitive digital PCR for SHIV-specific SIV *gag* gene regions. All of the SHIV-Ag85B- and SHIV-NI-inoculated macaques were euthanized and taken to necropsy from 264 weeks after the challenge, and proviral DNA analyses were performed on necropsy lymphoid tissue samples (Fig. 7C). Two SHIV-Ag85B-undetected macaques (#508096 and #912080) and all of the SHIV-NI-infected macaques showed plasma viremia at necropsy, and most of the infected macaques showed proviral DNA loads in the lymphoid tissues examined. In the SHIV-Ag85B-undetected macaques (#501006, #509106, #803032 and #901007), proviral DNA was not detected in any of the lymphoid tissues analyzed (Fig. 7C).

## Discussion

In the present study, we examined immune responses in cynomolgus macaques inoculated with SHIV-Ag85B and also the SHIV-specific immunity induced by an adjuvant molecule-expressing virus, SHIV-Ag85B, in the macaques after challenge with heterologous pathogenic SHIV89.6P. For this purpose, we genetically engineered a *nef*-deleted attenuated SHIV to harbor the Ag85B gene, so that Ag85B is produced locally when viral replication occurs. The SHIV-Ag85B-inoculated macaques showed a larger number of Gag/pol-specific IFN-γ-producing cells than that in SHIV-NI-inoculated macaques at 2 weeks after inoculation. Similarly, the SHIV-Ag85B-inoculated macaques showed a larger percentage of antigen-specific CD8^+^ T cells than that in the SHIV-NI-inoculated macaques. In a mouse model, co-injection of Ag85B plasmids with HIV Env DNA vaccine enhanced antigen-specific cellular immune responses (18). Our results suggest that the co-expression of a viral antigen and Ag85B stimulates antigen-specific cellular immune responses, which may help to increase the antiviral immune responses.

When we used cell lines, the replication of SHIV-Ag85B and that of parental SHIV-NI were similar. However, in the animal study, the viral loads in SHIV-Ag85B-inoculated macaques were lower than those in SHIV-NI-inoculated macaques. Moreover, proviral DNAs in PBMCs of SHIV-Ag85B-inoculated macaques were not detected at the early stage of infection. Similar results were obtained in the second experiment. In contrast, proviral DNAs in PBMCs of SHIV-NI-inoculated macaques were detected during the observation period. Our study also showed that inoculation of SHIV-Ag85B induced a remarkable antigen-specific T cell response against acute infection. One possible explanation is that the combination of Ag85B and a viral antigen mainly induced cellular immunity, as indicated by antigen-specific T cell responses, resulting in a more efficient immunomodulating effect against pathogenic SHIV. Induction of strong SHIV-specific cell-mediated immunity was also thought to be responsible for early virus eradication.

It is known that immune responses induced by an attenuated virus can provide protective immunity to a pathogenic virus. It has been reported that live attenuated SIV or SHIV vaccines can protect macaques against pathogenic SIV or SHIV challenges (8-10, 25, 26). In this study, all of the macaques inoculated with SHIV-Ag85B did not show any plasma viral load and proviral DNA at 8 weeks after injection of the virus. Moreover, most of the SHIV-Ag85B-undetected macaques, including those in the second experiment, did not show any evidence of SHIV89.6P infection at 20 weeks after the challenge with pathogenic SHIV89.6P (Fig. 3A). It should be mentioned here that PBMCs isolated from SHIV-Ag85B-inoculated macaques showed robust IFN-γ ELISPOT responses in the acute phase of pathogenic SHIV challenge. Moreover, the SHIV-Ag85B-undetected macaques showed increases in the percentages of Gag/pol-specific monofunctional and polyfunctional CD8^+^ T cells at 2 weeks after the challenge. In contrast, macaques in the group that received SHIV-NI did not effectively induce protective immunity. A SHIV genetically engineered to express the *Ag85B* gene enhanced the anamnestic cellular immune responses against a heterologous pathogenic SHIV89.6P challenge as well as the primary immune responses.

Innate immune responses have been thought to play an important role in both the control and pathogenesis of HIV or SIV infection (27-30). Furthermore, for the generation of adaptive immune responses, induction of innate immunity is crucial for vaccines to elicit potent antigen-specific immune responses. Attempts have been made to use various types of adjuvants for enhancing an immune response to an antigen, including vaccines or therapeutic cure against HIV. In fact, poxvirus- and adenovirus-based vaccines required the addition of an adjuvant to induce effective immune responses (6, 31, 32). For the generation of adaptive immune responses, induction of innate immunity is crucial for vaccines to elicit potent Ag-specific immune responses. As a result, it was demonstrated that SHIV-Ag85B might have a potent adjuvant activity as dsRNA recognized by the RIG-I receptor and enhanced not only local innate immunity but also systemic adaptive immunity.

Indian rhesus macaque AIDS models possessing MHC-I alleles, such as *Mamu-A*\*01, *Mamu-B*\*08, and *Mamu-B*\*17, tend to show spontaneous viral control after SIVmac challenge (33-36). Therefore, analysis of the MHC allele is important for HIV vaccine research using AIDS macaque models. In a previous study, we established an AIDS model using cynomolgus macaques of Asian origin that exhibited MHC genetic diversity populations. Moreover, these MHC-I alleles showed more genetic diversity than those of Mauritian cynomolgus macaques(37, 38). We analyzed the MHC class I alleles of the macaques used in this study. The macaques had variable MHC class I alleles and we did not find restrictive MHC class I alleles in the animals used in this study that are similar to *Mamu-A*\*01, *Mamu-B*\*08, and *Mamu-B*\*17 in rhesus macaques (data not shown). We excluded macaques with MHC alleles that are known to confer enhanced SHIV control. We then used a stringent animal model involving a pathogenic SHIV.

Approximately 50% of the macaques vaccinated with a cytomegalovirus-based SIV vaccine manifested complete control of viral replication shortly after SIVmac239 infection (39-41). Recently, it has been reported that “Miami Monkey” achieved durable HIV remission through adeno-associated virus-based delivery of broadly neutralizing antibodies against SHIV (42). In this study, the pathogenic virus was not detected in 57% of the macaques that had been injected with SHIV-Ag85B. Those macaques showed no detectable pathogenic virus measured as viral DNA in PBMCs and tissues or by adoptive transfer experiments and subsequent depletion of CD8+ cells. The adjuvant may provide the possibility of not only a preventive vaccine but also a functional cure of AIDS virus infection.

Adoptive transfer and CD8 depletion studies are useful for assessing residual replication-competent AIDS virus (41, 43, 44). The four SHIV-Ag85B-inoculated animals did not show viral RNA and provirus DNA in adoptive transfer and CD8^+^ cell depletion experiments. In previous studies, some live attenuated SIV vaccines were found to confer protection against SIVmac challenge. However, levels of protection were variable. Also, those attenuated vaccines could not completely protect against or eliminate SIV (13, 14, 25, 41). Therefore, we could not exclude the possibility that exceedingly low levels of replication-competent virus may still exist in the macaques. However, there was a clear difference between the macaques that exhibited viral replication and those that did not in adoptive transfer and CD8^+^ cell depletion experiments.

In summary, our results showed that immunization with SHIV-Ag85B provided completely protection against pathogenic SHIV by sterile immune responses. In SHIV-Ag85B-inoculated macaques, SHIV antigen-specific CD8^+^ T cell responses with polyfunctionality were rapidly induced. Surprisingly, four of the SHIV-Ag85B-inoculated macaques did not show virus replication in the adoptive transfer experiment and CD8^+^ cell depletion study. These results suggest that SHIV-Ag85B elicited viral antigen-specific CD8^+^ T cell responses against pathogenic SHIV and provide the possibility of eradicating a pathogenic lentivirus from infected cells. The results of this study provide further insights into the containment of HIV infections as well as new opportunities to develop a better therapeutic cure and vaccines.

## Materials and Methods

### Construction of SHIV-Ag85B

A recombinant SHIV was constructed according to the method reported previously (15, 16). The SHIV-nef vector, designated as SHIV-NI (Fig. S1A), was constructed from an infectious molecular clone of SHIV-NM3rN (45). In SHIV-NI, the *nef* gene was replaced by some unique restriction enzyme sites including ClaI and ApaI sites. The Ag85B gene was amplified by PCR from *Mycobacterium kansasii* as a template using 5′-ATATCGATACCATGTTCTCCCGTCCCGGGCT-3′ (ClaI) and 5′-TAGGGCCCCTA GCGGGCGCCCAGGCTGG-3′ (ApaI) primers. The PCR product was then digested with the restriction enzymes for ClaI and ApaI sites. This plasmid was designated as pSHIV-Ag85B. SHIV-Ag85B was prepared by transfecting pSHIV-Ag85B into 293T cells using FuGENE 6 Transfection Reagent (Roche Diagnostics, Indianapolis, IN), and the culture supernatant was stored at 48 h after transfection in liquid nitrogen until use.

### Virus stocks

SHIV-Ag85B, SHIV-NI and SHIV89.6P were used in this study. These viral stocks were propagated in PBMCs from cynomolgus macaques. Briefly, PBMCs were separated by a standard Ficoll density gradient separation method and cultured in RPMI1640 supplemented with 10% fetal bovine serum, 2 mM L-glutamine, and 100 units/ml of IL-2 (Shionogi) and then stimulated with phytohemagglutinin for 72 h. The cells were infected with SHIV-Ag85B, SHIV-NI or SHIV89.6P at a multiplicity of infection (MOI) of 0.1. Half of the culture medium was replaced with a fresh culture medium every three days, and cell-free supernatants were collected between 6 and 9 days after infection. The tissue culture 50% infectious dose (TCID_50_) of each of the SHIVs was measured using M8166 cells. The TCID_50_ values of viral stocks were 5 × 10^4^ for SHIV-Ag85B, 4.7 × 10^4^ for SHIV-NI and 3 × 10^5^ for SHIV89.6P.

### Detection of Ag85B protein

M8166 cells were infected with SHIV-Ag85B at an MOI of 0.1 and incubated for 1 h. The cells were washed three times with PBS and then incubated for another 48 h in the culture medium. After three further washings with PBS, the cells were lysed in PBS containing 1.5 M urea, 2% NP-40 and 5% 2-mercaptoethanol and then separated by SDS-PAGE, transferred by electroblotting onto a nitrocellulose membrane, and blocked with 5% nonfat dry milk in PBS containing 0.01% Tween 20 (PBST). Following three washings with PBST, the membrane was incubated with a rabbit anti-Ag85B polyclonal antibody for 2 h. The membrane was washed three times with NBT/BCIP (Roche Diagnostics, Mannheim, Germany) before being incubated with alkaline phosphatase-labeled anti-rabbit IgG (New England Biolabs, Beverly, MA).

### Virus replication of human and macaque cells

To investigate the kinetics of virus replication, CEM×174 were infected with SHIV-Ag85B and SHIV-NI at an MOI of 0.01 and incubated for 1 h. Half of the culture supernatant was harvested with subsequent addition of a new medium every 3 days. Virus replication kinetics was monitored by SIV Gag p27 production in the supernatant using an SIV core antigen ELISA assay kit (Beckman Coulter, Miami, FL). To assess the properties of SHIV-Ag85B, macaque (HSC-F) cell lines were also inoculated with the virus.

### Innate immune responses

CEM×174 cells (2 × 10^5^ cells per well) were infected with SHIV-Ag85B or SHIV-NI at an MOI of 0.2 and cultured for 48 hours. The cells were harvested, and mRNA was extracted using an RNeasy Mini Kit (QIAGEN) and then reverse-transcribed to cDNAs using an Omuniscript system (QIAGEN). The cDNA was subjected to real-time PCR for IFN-α, IFN-β, IFN-γ, TNFα, Il6, RIG-I, MDA5, LGP2, and β-actin selectin using a LightCycler (Roche Applied Science, Tokyo, Japan). The specific primers for each target (listed below) and probes were designed by Universal ProbeLibrary Assay Design Center (Roche Applied Science). Primers used in this study were 5′-CCCTCTCTTTATCAACAAACTTGC-3′ and 5′-TTGTTTTCATGTTGGACCAGA-3′ for IFN-α, 5′-CGACACTGTTCGTGTTGTCA-3′ and 5′-GAAGCACAACAGGAGGAGCAA-3′ for IFN-β, 5′-GGCATTTTGAAGAATTGGAAAG-3′ and 5′-TTTGGATGCTCTGGTCATCTT-3′ for IFN-γ, 5′-GATGAGTACAAAAGTCCTGATCCA-3′ and 5′-CTGCAGCCACTGGTTCTGT-3′ for IL6, 5′-AGCCCATGTTGTAGCAAACC -3′ and 5′-TCTCAGCTCCACGCCATT-3′ for TNFα, 5′-TGGACCCTACCTACATCCTGA-3′ and 5′-GGCCCTTGTTGTTTTTCTCA-3′ for RIG-I, 5′-AGGCACCATGGGAAGTGAT-3′ and 5′-GGTAAGGCCTGAGCTGGAG-3′ for MDA5, 5′-ATGTGAACCCCAACTTCTCG-3′ and 5′-GCTTCCAGTCCTTGAAGACTTT-3′ for LGP2, and 5′-CCAACCGCGAGAAGATGA-3′ and 5′-CCAGAGGCGTACAGGGATAG-3′ for β-actin.

### Animals

We used 22 adult cynomolgus macaques (from Indonesia, Philippines, and Malaysia), all of which were negative for SIV, simian type D retrovirus, simian T-cell lymphotropic virus, simian foamy virus, Epstein-Barr virus, cytomegalovirus, and B virus. Seven cynomolgus macaques were intravenously inoculated with 10^4^ TCID_50_ of SHIV-Ag85B, five macaques were intravenously inoculated with 10^4^ TCID_50_ of SHIV-NI and another five macaques were intravenously inoculated with 10^4^ TCID_50_ of SHIV89.6P. To evaluate viral replication, a further 7 macaques were inoculated intravenously with SHIV-Ag85B (second experiment; Exp 2). To evaluate the protective efficacy of SHIV-Ag85B or SHIV-NI in inoculated macaques, the macaques were challenged intravenously with heterologous pathogenic SHIV89.6P at 37 weeks after inoculation. As controls, a further 5 naïve macaques were challenged intravenously with SHIV89.6P. The design of the macaque study is outlined in Fig.1A.

### Adoptive transfer experiment

For the adoptive transfer experiment, 15 ml of blood and 1 to 3 × 10^7^ cells of lymphoid tissues from SHIV-Ag85B or SHIV-NI-inoculated macaques, collected at 212 weeks after SHIV89.6P challenge, were infused intravenously into healthy macaques. Viral and proviral DNA loads were assessed in recipient macaques during the observation period after adoptive transfer.

### CD8 depletion experiment

For the CD8^+^ cell depletion experiment, macaques received a single intravenous infusion of 10 mg/kg of a CD8α CDR-grafted rhesus IgG1 antibody (M-T807) (NIH Nonhuman Primate Reagent Resource). Antibody cTM-T807 was intravenously inoculated at 252 weeks after the challenge. CD8^+^ T cell counts and viral and proviral DNA loads were assessed in recipient macaques during the observation period after the CD8^+^ depletion experiment.

### Sample collection

Blood was collected periodically using sodium citrate as an anticoagulant and used for determination of CD4^+^ T cell counts, quantification of plasma viral loads, and immunological analysis. Lymphoid tissue samples were obtained by biopsy at 34 weeks after virus infection and were used for determination of proviral DNA loads and histopathology. These studies were performed in TPRC, National Institutes of Biomedical Innovation, Health and Nutrition (NIBIOHN) after approval by the Committee on the Ethics of Animal Experiments of NIBIOHN in accordance with the guidelines for animal experiments at NIBIOHN. The animals were used under the supervision of the veterinarians in charge of the animal facility. The endpoint for euthanasia was determined by typical clinical symptoms of AIDS such as weight loss, diarrhea, and neurologic syndrome. These diagnostics were performed by the veterinarians.

### Preparation of DNA samples and amplification of the SHIV *gag* gene by nested PCR

For determining the proviral DNAs in SHIV-Ag85B-inoculated macaques, nested PCR was used to amplify a fragment of the *gag* gene segment. Proviral DNA was extracted from PBMCs of the inoculated macaques. Cellular DNAs were extracted using DNeasy tissue kits (QIAGEN). Nested PCR was performed using TaKaRa Ex Taq (Takara Bio Inc., Shiga, Japan). The initial and nested PCR protocols have been described elsewhere(46, 47). Primers used in this study were Outer SIV*gag*-F (5’-CCATTAGTGCCAACAGGCTCAG-3’) and Outer SIV*gag*-R (5’-CCCCAGTTGGATCCATCTCCTG-3’) for first-round PCR and Nested SIV*gag*-F (5’-ACTGTCTGCGTCATCTGGTG-3’) and Nested SIV*gag* –R (5’-GTCCCAATCTGCAGCCTCCTC-3’) for second-round PCR. After the second amplification, 10 μl of the nested PCR-amplified product was run on 1.0% agarose gel, and DNA bands were visualized by staining with ethidium bromide. The lowest concentration of plasmid SIV DNA that could be detected with this PCR method in the first amplification with the outer gag primer pair was 103 copies. Upon further amplification with nested/internal gag primers, a single copy of plasmid DNA could be routinely detected (46, 47).

### Stability of the inserted *Ag85B* gene *in vivo*

Proviral DNA was extracted from 1 × 10^6^ PBMCs of the inoculated macaques. When the virus was re-isolated, CD8^+^-depleted PBMCs, which were co-cultured with M8166 cells, were also monitored. Cellular DNAs were extracted using DNeasy tissue kits (QIAGEN). To check the stability of the inserted *Ag85B* gene in SHIV-Ag85B, the proviral DNA fragments covering the inserted *Ag85B* gene in SHIV-Ag85B were amplified by PCR with primers. The primer sequences are shown above.

### Plasma viral RNA loads

The levels of SHIV infection were monitored by measuring plasma viral RNA loads using highly sensitive quantitative real-time RT-PCR as described previously (37, 48). Briefly, viral RNA was isolated from plasma using a MagNA PureCompact Nucleic Acid Isolation kit (Roche Diagnostics). Real-time RT-PCR was performed using a QuantiTec Probe RT-PCR kit (Qiagen) and a LightCycler 480 thermocycler (Roche Diagnostics, Rotkreuz, Switzerland). The SIVmac239 *gag* gene was amplified with the probe 5′-FAM-TGTCCACCTGCCATTAAGTCCCGA-TAMRA-3′ (where FAM is 6-carboxyfluorescein and TAMRA is 6-carboxytetramethylrhodamine) and the primers 5′-GCAGAGGAGGAAATTACCCAGTAC-3′ and 5′-CAATTTTACCCAGGCATTTAATGTT-3′. The limit of detection was calculated to be 100 viral RNA copies per ml.

### Proviral DNA loads

A DNA sample was extracted from PBMCs and lymphoid tissues using a DNeasy tissue kit (QIAGEN) according to the manufacturer’s protocol. Ultrasensitive digital PCR was performed in the QX200 Droplet Digital PCR system (Bio-Rad). Twenty μl of a reaction mixture containing 2 μl of a DNA sample, ddPCR supermix for probes (no dUTP) (Bio-Rad), 900 nM of each primer, 200 nM of the probe, and demineralized water was prepared. The mixture was placed into the DG8 cartridge with 70 μl of droplet generation oil (Bio-Rad), and the droplets were formed in the droplet generator (Bio-Rad). Subsequently, the droplets were transferred to a 96-well microplate. PCR amplification was performed using the following program: initial denaturation and stabilization at 95°C for 10 min, 40 cycles of denaturation at 94°C for 30 s, and annealing/extension at 57°C for 60 s, followed by 10 min at 98°C. Subsequently, droplets were sorted and analyzed in a QX200 droplet reader (Bio-Rad) using the software QuantaSoft v1.6 (Bio-Rad). Samples were only considered if more than 20000 droplets were read. The cell numbers were monitored by highly sensitive quantitative real-time PCR as described previously (49). DNA samples were extracted from PBMCs and lymphoid tissues using a DNeasy tissue kit (QIAGEN) according to the manufacturer’s protocol. The cell number was validated by detection of the cellular interleukin-4 (IL-4) sequence with the rhesus IL-4-specific primers 5′-TGTGCTCCGGCAGTTCTACA-3′ and 5′-CCGTTTCAGGAATCGGATCA-3′ and the probe 5′-FAM-TGCACAGCAGTTCCACAGGCACAAG-TAMRA-3′.

### CD4+ and CD8+ T-cell counts

One hundred microliters of whole blood from each of the cynomolgus macaques was stained with combinations of fluorescence-conjugated monoclonal antibodies: anti-CD3 (clone SP34-2, Alexa700; BD), anti-CD4 (clone L200, PerCP-Cy5.5; BD), and anti-CD8 (clone DK25, APC; Dako). Flow cytometry was performed on a FACSCanto II flow cytometer (BD). The data were analyzed using FACSDiVa software.

### Immunohistochemistry

Lymphoid tissue samples were fixed in 4% paraformaldehyde in PBS at 4 °C overnight and embedded in paraffin wax. Sections were deparaffinized by pretreatment with 0.5% H_2_O_2_ in methanol and then subjected to antigen retrieval with target retrieval solution (Dako S1700, pH 6.1) followed by heating in an autoclave for 20 min at 121 °C. The sections were then incubated with an anti-human CD4 mouse monoclonal antibody (1:30; NCL-CD4; Novocastra Laboratories Ltd., United Kingdom) or an anti-SIV mouse monoclonal antibody (1:100; SIV-Nef; Santa Cruz Biotechnology) at 4 °C for 24 h. Following brief washes with a buffer, the sections were incubated with the EnVision™+ Dual Link-HRP system (Dako) as a secondary stage for 60 min. Labeling was “visualized” by treating the sections with chromogen 3,3’-diaminobenzidine tetroxide (Dojin Kagaku, Japan) and H_2_O_2_. The sections were then counterstained with haematoxylin.

### Analysis of polyfunctional Gag/pol-specific T-cell responses

Antigen-specific cellular immune responses were assessed in multiparameter intracellular cytokine staining (ICS) assays as described previously (50, 51). Briefly, thawed cryopreserved PBMCs were co-cultured for 6 h with autologous herpes virus papio-immortalized B-LCL cells that had been infected with a recombinant vaccinia virus expressing SIV Gag/pol. They were incubated with brefeldin A (BD) for the final 5 h of stimulation. Then immunostaining was performed using a CytofixCytoperm kit (BD) and the following monoclonal antibodies: anti-CD3 (clone SP34-2, Alexa700; BD), anti-CD4 (clone L200, APC-H7; BD), anti-CD8 (clone DK25, APC; Dako), anti-IFN-γ (clone 4S.BS, PE; BD), anti-TNF-α (clone MAM11 PE-Cy7), and anti-IL-2 (clone MQ-1-17H12). A fixable-dead-cells stain kit (Invitrogen) was used to exclude dead cells from the analysis. Samples were fixed with 1% freshly prepared paraformaldehyde for at least 1 h and then analyzed in a FACSCanto II flow cytometer (BD). Data were analyzed using FACSDiVa (BD) software.

### Antigen-specific IFN-γ ELISPOT assay

The number of antigen-specific IFN-γ-producing cells in PBMCs was determined by ELISPOT analysis as described previously (37). Ninety-six-well, flat-bottom plates were coated with an anti-IFN-**γ** monoclonal antibody (clone MD-1; U-Cytech, Utrecht, Netherlands) and blocked with 2% BSA in PBS. Briefly, thawed cryopreserved PBMCs were co-cultured for 9 h with autologous herpes virus papio-immortalized B-LCL cells that had been infected with a recombinant vaccinia virus expressing SIV Gag/pol or Ag85B. Gold-labeled anti-biotin IgG solution (U-Cytech, Utrecht, Netherlands) was added to the washed plates and then incubated for 1 h at 37 °C. Spot-forming cells (SFCs) were counted using the KS ELISPOT compact system (Zeiss) after a 15-min reaction with an activator mix (U-Cytech, Utrecht, Netherlands). An SFC was defined as a large black spot with a fuzzy border.

### Memory phenotype of SHIV-specific CD8^+^ T cells

PBMCs treated as described above were immunostained using a CytofixCytoperm kit (BD) and the following monoclonal antibodies: anti-CD3 (clone SP34-2, Alexa700; BD), anti-CD4 (clone L200, APC-H7; BD), CD8 (clone DK25, APC; Dako), CD28 (clone CD28.2, ECD; Beckman-Coulter), CD95 (clone DX2, PE-Cy7; BD), and IFN-γ (clone 4S.BS, PE; BD). A fixable-dead-cells stain kit (Invitrogen) was used to exclude dead cells from the analysis. The percentages of CD8^+^ T cells in CM and EM populations were determined by using a combination of CD28 and CD95 markers. Samples were fixed with 1% of freshly prepared paraformaldehyde for at least 1 hour and then analyzed using a FACSCanto II flow cytometer (BD). The data analysis was conducted using FACSDiVa (BD) software.

### Virus-specific antibody titer

Anti-SHIV titers were determined by using a commercial passive agglutination method (Serodia-HIV1/2; Fuji rebio, Japan). Isolated plasma samples were serially diluted and assayed. The antibody titer was measured by fourfold serial dilution of each sample, as recommended by the manufacturer.

### Statistical analysis

Data are represented as means ± SD. A *P* value of <0.05 is considered significant. Statistical analyses were performed using the Mann-Whitney *U* test. Statistically significant differences compared with the control are indicated by asterisks.

## Acknowledgements

We thank K. Ueda for excellent technical assistance. Special thanks go to the members and veterinary staff of HAMRI CO., LTD, particularly J. Sawata, T. Horikawa, M. Kogure, S. Yokota, and H. Ayukawa for their technical expertise and assistance with animal care. We also thank the members of Corporation for Production and Research of Laboratory Primates, especially K. Hanari, N. Nakano, and K Ohto for their support in animal experiments. This work was partially supported by Health Science Research Grants from the Ministry of Health, Labor and Welfare of Japan and the Ministry of Education, Culture, Sports, Science and Technology, Japan, and the Japan Agency for Medical Research and Development, AMED (19ak0101047h0004, 19fk0410025h0001, 19ak0101068h0003 and 19fk0410011h0702). The funders had no role in study design, data collection and analysis, decision to publish, or preparation of the manuscript.

Y.Y. and T.O. designed the experiments. T.O., Y.S., and K.M. constructed and validated the SHIVs used in this study. T.O., T.K., and T.Y. performed the virologic assays. T.O., Y.T., and M.N.A. perfomed the immunologic assays. T.O performed tissue analysis. T.O., K.M., T.Y., and Y.Y. wrote the manuscript.

The authors declare no competing interests.

